# Parallel genomic basis of age at maturity across spatial scales in Atlantic Salmon

**DOI:** 10.1101/2022.09.09.507321

**Authors:** Tony Kess, Sarah J. Lehnert, Paul Bentzen, Steven Duffy, Amber Messmer, J. Brian Dempson, Jason Newport, Christopher Whidden, Martha J. Robertson, Gerald Chaput, Cindy Breau, Julien April, Carole-Anne Gillis, Matthew Kent, Cameron M. Nugent, Ian R. Bradbury

## Abstract

Complex traits often exhibit complex underlying genetic architectures resulting from a combination of evolution from standing variation, hard and soft sweeps, and alleles of varying effect size. Increasingly, studies implicate both large-effect loci and polygenic patterns underpinning adaptation, but the extent that common genetic architectures are utilized during repeated adaptation is not well understood. Sea age or age at maturation represents a significant life history trait in Atlantic Salmon *(Salmo salar)*, the genetic basis of which has been studied extensively in European Atlantic populations, with repeated identification of large-effect loci. However, the genetic basis of sea age within North American Atlantic Salmon populations remains unclear, as does the potential for a parallel trans-Atlantic genomic basis to sea age. Here, we used a large SNP array and low coverage whole genome re-sequencing to explore the genomic basis of sea age variation in North American Atlantic Salmon. We found significant associations at the gene and SNP level with large-effect loci (*vgll3, six6)* previously identified in European populations, indicating genetic parallelism, but found that this pattern varied based on both sex and geographic region. We also identified largely non-repeated sweep signatures and sets of highly predictive loci associated with sea age among populations and sexes within North America, indicating polygenicity and low rates of repeated genomic parallelism. Despite low genome-wide parallelism, we uncovered a set of conserved molecular pathways associated with sea age that were consistently enriched among comparisons. Together, our results indicate parallelism of the molecular basis of sea age in North American Atlantic Salmon across large-effect genes and molecular pathways despite population-specific patterns of polygenicity. These findings reveal roles for both contingency and repeated adaptation at the molecular level in the evolution of life history variation.

## Introduction

Understanding the genomic basis of adaptive traits is important for uncovering the molecular mechanisms that generate biodiversity (Hoban et al., 2016). A key component of understanding adaptive genetic variation is in identifying the predictability of genomic patterns underlying repeated adaptation, providing insight into the number of molecular solutions for ecological challenges (Blount et al., 2018; Elmer and Meyer 2011). Theoretical and empirical studies have indicated that variability among genetic architectures underlying adaptive traits may have consequences for their genomic parallelism (Bolnick et al., 2018; Yeaman et al., 2018). Parallelism at the genomic level is expected to occur with the greatest frequency in scenarios of shared ecological conditions and selection (Kaeuffer et al., 2012), phylogenetic similarity (Conte et al., 2012), shared standing variation (Ralph & Coop 2015), and large-effect loci controlling a large proportion of phenotypic variance (Yeaman 2015). However, many traits exhibit polygenic architecture, and models of polygenic adaptation indicate small changes in allele frequency across many genetic pathways could reduce genomic parallelism (Barghi et al., 2020; Fagny & Austerlitz 2020; Yeaman 2015). Large-scale follow-up studies on the genetic basis of adaptive traits with previously identified large-effect loci have also identified polygenic patterns explaining additional variation (Kreiner et al., 2020, Sinclair-Waters et al., 2020), suggesting polygenicity may be common. Currently, the extent that simple and repeatable or polygenic and variable genetic architectures are more frequent during repeated adaptation remains unknown, requiring further study in the wild to characterize the genomic parallelism of adaptive traits.

Salmonids exhibit an extensive array of adaptive diversity, with varying rates of underlying genomic parallelism (Jeffery et al., 2017; Klemetsen et al., 2003; Salisbury et al., 2022). Atlantic Salmon (*Salmo salar*) is a culturally, ecologically, and economically significant species with an anadromous life cycle consisting of fresh water residency followed by a period at sea with enhanced growth rates prior to sexual maturation upon returning to fresh water to spawn (Mobley et al., 2021). Sea age at first maturation varies both among individuals (Jonsson & Jonsson, 2007) and rivers (Hutchings & Jones, 1998) and represents a variable life history strategy based on investment in reproduction timing and fecundity (Garant et al., 2003), with both genetic (Johnston et al., 2014) and environmental underpinnings (Friedland et al., 2000). Atlantic Salmon is also iteroparous and can spawn multiple times; sea age at first maturation age is demarcated by the number of winters spent at sea, also known as “sea age” or “sea winters”, with fish maturing after a single sea winter commonly known as grilse or one-sea-winter salmon (1SW) (Johnston et al., 2014) and fish first maturing after two winters at sea (2SW), three winters at sea (3SW) and older commonly referred to as multi-sea-winter (MSW) salmon. Declines in Atlantic Salmon stocks across both Europe and North America (Lehnert et al., 2019, Olmos et al., 2020), and reductions in the number of multi-sea-winter (MSW) salmon across years (Olmos et al. 2020) suggest recent selection against older age at first maturation (Czorlich et al., 2018), and highlights the need for greater understanding of this trait to inform conservation of life history variation within this species.

The genetic basis of sea age has been studied primarily in European populations (Barson et al., 2015; Sinclair-Waters et al., 2021), but the degree of genomic parallelism with North American populations remains unclear (e.g., Boulding et al., 2019). Genomic investigation of this trait in rivers in Norway identified > 30% of sea age variation controlled by a large-effect locus, (*vgll3*, Barson et al., 2015) also associated with adiposity regulation and puberty onset in humans (Cousminer et al., 2013; Perry et al., 2014). However, significant associations with major effect loci in North American populations have remained elusive, indicating a complex genetic architecture (Boulding et al., 2019; Kusche et al., 2017; Mohamed et al., 2019). Divergence between European and North American Atlantic Salmon populations is high (*F*_ST_ =0.26) and genome-wide (Lehnert et al., 2020), reflecting extensive periods of isolation (∼600,000 years) for the potential evolution of distinct genetic architectures of maturation. However, secondary contact between North American and European lineages during deglaciation has also been identified, (Rougemont & Bernatchez 2018) and has provided the opportunity for introgression of adaptive variation (Lehnert et al., 2019), potentially leading to a shared genetic basis of maturation at least in some regions. Genome-scale investigations into sea age variation in wild Atlantic Salmon populations within North America have not yet been carried out and have the potential to reveal the genetic architecture and parallelism of this important life history trait.

Here, we used low-coverage whole genome re-sequencing (WGS) and a single nucleotide polymorphism (SNP) array and individuals from rivers across eastern North America to carry out genome wide association (GWA) and genome scans for variation associated with sea age in Atlantic Salmon. To detect polygenic patterns associated with sea age, we then conducted tests of incomplete selective sweeps and machine-learning prediction of individual sea age from a panel of genome-wide markers. Finally, we used gene set enrichment to explore the biological and molecular function of loci exhibiting polygenic associations with sea age variation. Our findings here provide insight into factors driving parallelism of individual genes and higher-level conserved pathways underlying traits with variable and polygenic backgrounds.

## Methods

### Sampling and genotyping: SNP array

For site-level genomic analysis of population structure and sea age association, we used new and previously genotyped samples from published sources (Lehnert et al., 2020; Wringe et al., 2018), using a previously developed Atlantic Salmon 220K SNP Axiom Array (Barson et al., 2015). We used genotype data at 97,566 SNPs that passed Axiom quality filters, with minor allele frequencies (maf) greater than 0.05. For all analyses, genotype data were retained for 658 individuals from 26 sites with population-level mean proportion of maiden 1SW fish summarized from Canadian federal (Fisheries and Oceans Canada) and Quebec provincial (Ministère des Forêts, de la Faune et des Parcs) salmon counts from 1984-2020 (Figure 1A, Supplementary Table 1). Individual genotypes were exported in plink format (Chang et al., 2015), and subsequent data filtration and conversions for analysis of SNP Array data were carried out in *plink* 1.90b6.16.

**Figure 1.**
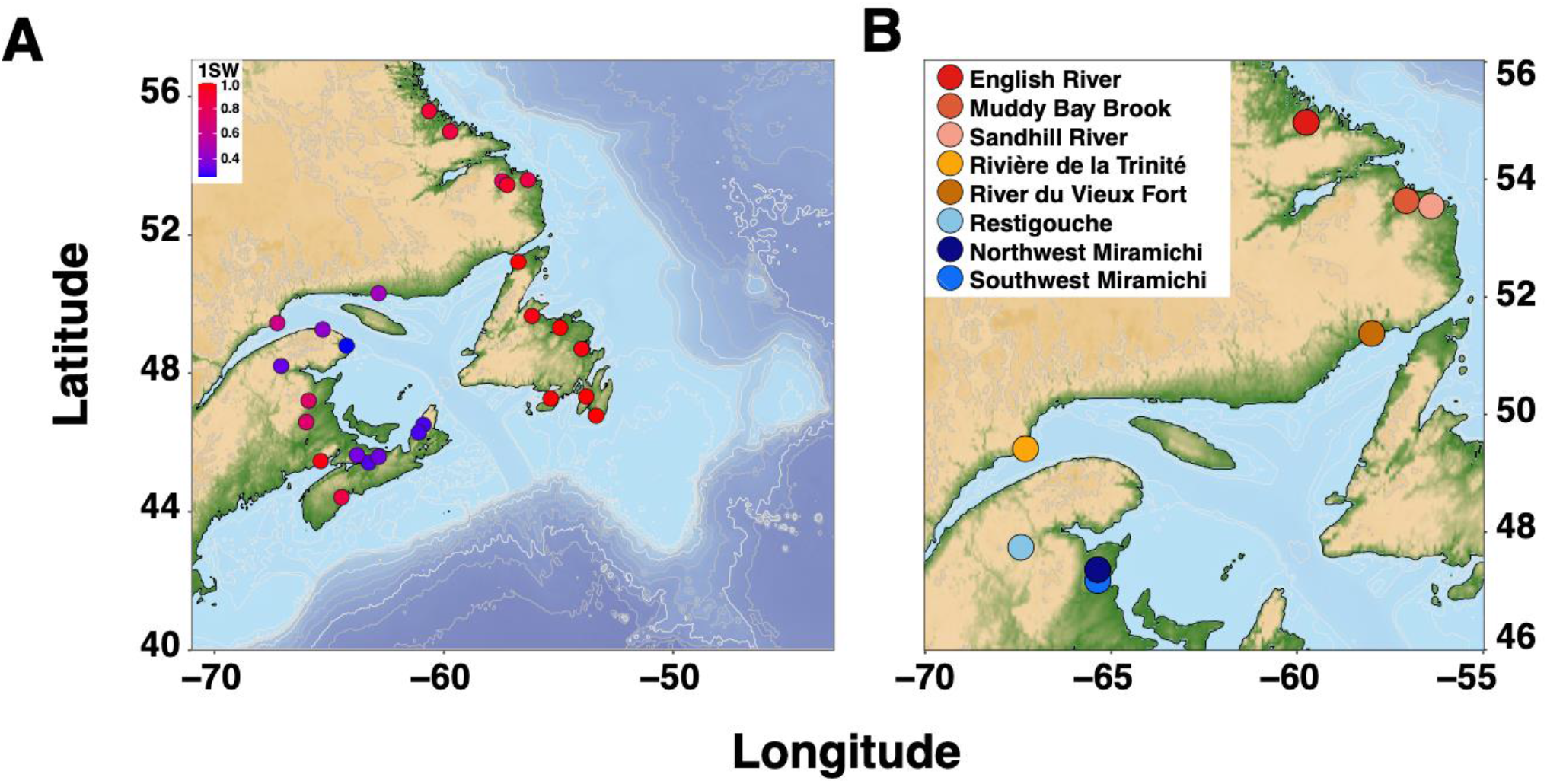
Map locations coloured by grilse proportion from 26 sampled rivers for individuals genotyped on the SNP array (A), and locations from 8 sampled rivers for samples with individual sea-age genotyped using low depth resequencing (B).

### Sampling and genotyping: WGS

To identify associations between genomic data, population structure and sea age variation at the individual level, we carried out low coverage whole genome re-sequencing as in Therkildsen & Palumbi (2017) of 582 male and female wild salmon from eight rivers in North America (Figure 1B, Supplementary Table 2). Atlantic Salmon were caught at fish counting fence traps or by angling fisheries, and were sampled for fin clips for DNA extraction, and scales to identify sea age. Sex was confirmed for each individual with PCR of the *Salmo salar* sex determining region (sdY). To prepare whole genome re-sequencing libraries, we extracted genomic DNA following Mayjonade et al., (2016), using Longmire’s buffer (Longmire et al., 1997) for tissue lysis. We then used a modified version of the protocol described in Therkildsen & Palumbi (2017) and scaled down reaction volumes of Nextera DNA Flex Library Prep Kits (Illumina) to 0.13x the volume of the standard Illumina protocol. For library amplification, we used Kapa Hi-Fidelity Library Amplification Kits (Roche) in 20 µL reactions with 4µL Nextera Unique Dual Indexes Set A (Illumina). We then quantified libraries by Qubit (ThermoFisher) and checked average fragment size using an Agilent Bioanalyzer. We normalized libraries from 96-well plates to equimolar concentrations prior to combining as a single pool per lane of sequencing, Libraries were then sequenced on six lanes of an Illumina NovaSeq6000 S4 at the Genome Quebec Centre d’Expertise et de Services.

Quality of sequenced libraries was checked using FastQC, (Andrews 2010). We used *cutadapt* 2.1 (Martin 2011) to remove the leading 15 bases, adapter content, bases falling below of q score of 10, and any read with less than 40 remaining base pairs. Trimmed reads from each individual were aligned to the 29 chromosomal contigs from the ICSASG_V2 *Salmo salar* reference assembly (GCA_002910315.2) using *bwa mem* 0.7.17 (Li 2013). Subsequent steps followed Best Practices recommendations for *Genomic Analysis Toolkit* (GATK 3.7, DePristo et al., 2011): we first removed duplicate reads using the *PicardTools* 2.20.6 *MarkDuplicates* function. De-duplicated reads were then sorted and realigned around potential insertions and deletions using *RealignerTargetCreator* and *IndelRealigner* functions in *GATK*.

Next, we estimated genotype likelihoods, as well as directly called genotypes for each individual using Analysis of Next Generation Sequencing Data (*ANGSD* 0.935, Korneliussen et al., 2014). We output genotype likelihoods for each chromosome as beagle format likelihood files (Browning & Browning 2007) for all SNPs passing quality filers (-*minMapQ* 30 -*minQ* 20 *-SNP_pval* 2e-6 *-uniqueOnly* 1 *-remove_bads* 1), and ensuring greater than 80% (-*minInd* 470) of individuals had genotypes and a minimum of 500 reads were present per locus (*-setMinDepth* 500). Genotype calls from each chromosome were exported as bcf files using the -*dobcf* flag, and converted to vcf files (Danecek et al., 2011) using *bcftools* 1.11 (Li et al., 2009). We then assessed mean depth of vcf files using *vcftools* 0.1.16, and then performed phasing and imputation of vcf files using *Beagle* 4.0 (Browning & Browning 2007), resulting in 9,895,443 SNPs with a mean coverage of 4.11x for downstream analysis.

### Population structure

We estimated population structure of Axiom Array data using principal component analysis (PCA) in the R package *pcadapt* (Privé et al., 2020), assuming K = 5 ancestral populations, based on examination of Scree plots (Supplementary Figure 1) and maf = 0.05. We also explored population structure using the sparse non-negative matrix factorization algorithm (*snmf*, Frichot et al., 2014) implemented in the *R* package *LEA* (Frichot & Francois 2015), with K=5 ancestral populations.

Population structure in WGS samples was quantified using genotype likelihoods in *PCAngsd* 1.02 (Meisner & Albrechtsen 2018). This method accommodates low coverage whole genome re-sequencing data through modelling genotype uncertainty in PCA for continuous estimation of population structure, and in non-negative matrix factorization for estimation of admixture components. To identify more fine-scale levels of population stratification for use as covariates in GWA, we repeated PCA separately on each regional grouping of samples, stratified by sex.

### PCA selection scan

Loci exhibiting signatures of selection associated with population structure were detected in SNP array data by estimating *p* values for each SNP using the Mahalanobis method in *pcadapt* with K = 5. Adjusted *p*-values for False Discovery Rate (FDR, Storey & Tibshirani 2003) were then estimated using the *qvalue* R packages (Storey 2015), and selected SNPs with *q* < 0.05 as exhibiting significant signatures of selection associated with population structure. To detect regions of elevated population-structure associated with divergence in WGS data, we estimated *p* values for each SNP from its loading on each PC axis, assuming a χ^2^ distribution of PC loadings, and then selected FDR-adjusted *q* values < 0.05.

### Genome-wide association with sea age

Genome-wide association with SNP array data was conducted using site-level grilse proportions, which reflect the average number of 1SWfish per sample river. Per-SNP associations with grilse proportion were estimated using latent factor mixed models with *LFMM 2* in *LEA* (Caye et al., 2019). We specified K=1 to test for association without correction for population structure, and K=2 to account for variation associated with the PC1 axis which separated individuals by site and grilse proportion (Figure 2B). To characterize multi-locus associations with grilse proportion, we conducted redundancy analysis (RDA) using the *vegan* R package (Oksanen et al., 2020), using grilse proportion to predict genome wide variation. Significance of the RDA model was tested using the *anova*.*cca* function with 999 permutations. Confounding of GWA scores by population structure was tested using variance partitioning with the *varpart* function to estimate the variation in grilse proportion explained by the first five principal components of genetic variation from PCA.

**Figure 2.**
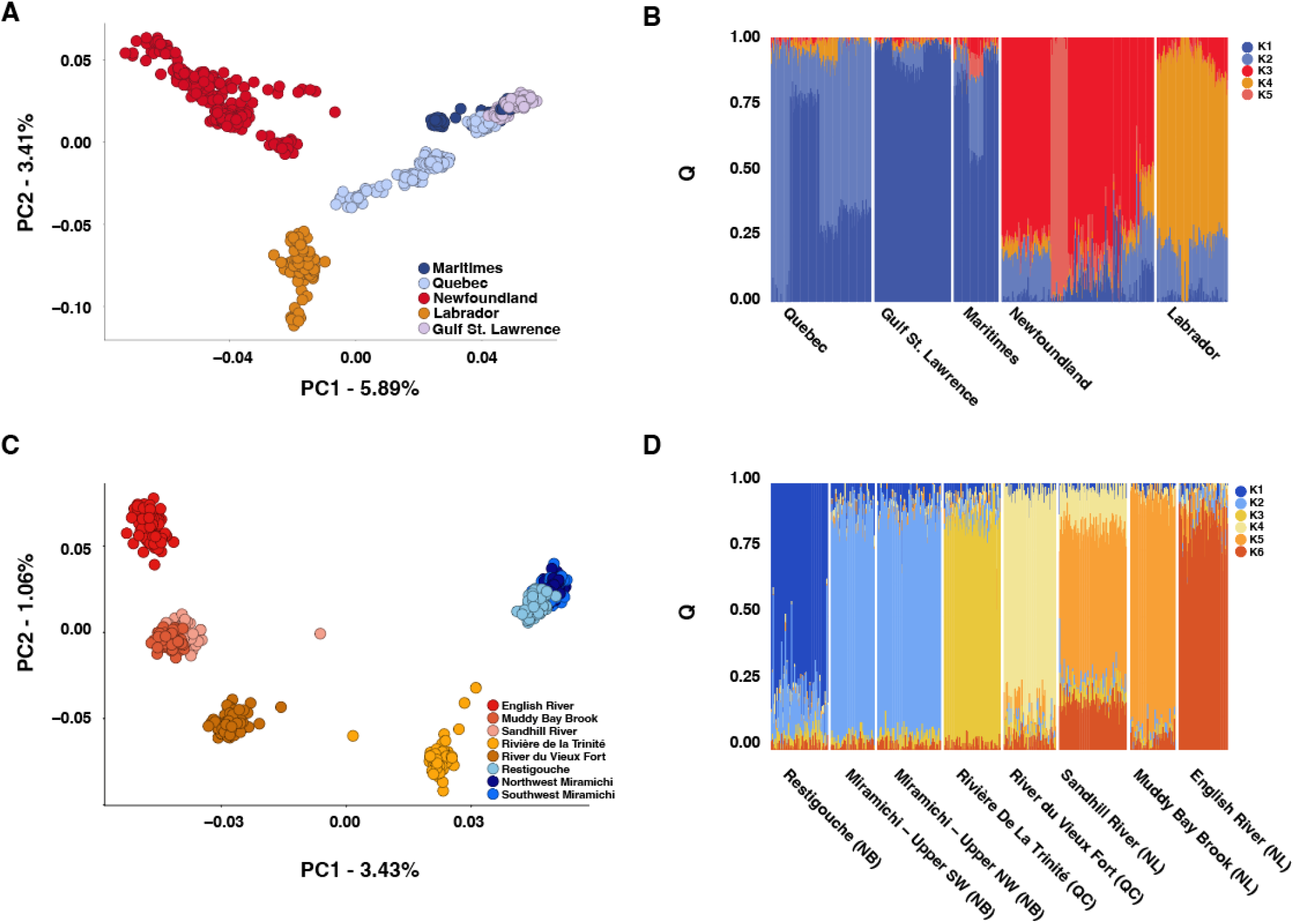
Population structure inferred from principal component (PC) analysis in *pcadapt* coloured by geographic origin (A), and individual admixture proportions identified from sparse non-negative matrix factorization in *LEA* of individuals genotyped on the SNP array (A), and population structure inferred from principal component analysis (C), and admixture proportions across regions identified from non-negative matrix factorization (D) in *PCAngsd* using WGS samples.

We used the association functions in *ANGSD* to test for association with individual sea age phenotypes in whole genome re-sequencing samples from genotype likelihoods using a latent genotype model (-*doAsso* 4, Jørsboe & Albrechtsen2022), to account for genotype uncertainty. Individual sea age was included as a binary phenotype, as all samples compared in this study exhibited a sea age at first maturity of one or two years, and only a small subset (n=11) exhibited total sea age greater than two, but also were assigned a sea age of one, indicating repeat spawning. Association analysis was carried out on genotype likelihoods from the phased and imputed vcf of all samples, without correction for population structure, as well as with the scores from the first two PCs as covariates. We then repeated this analysis within each regional grouping (within southern Quebec and New Brunswick rivers, and within northern rivers in Quebec and Labrador) and sex, with the first PC included in each analysis as a covariate to correct for population structure. Per-locus Weir & Cockherham’s *F*_ST_ (1984) was estimated with *vcftools* from phased vcf files between these same groups based on separation by sea age class, as a second measure of divergence associated with sea age.

### Detection of selection sweeps: WGS

We next explored signals of polygenic selection between groups in WGS data by estimating XP-nSL (Szpiech et al., 2021), an extension of the nSL haplotype-based statistic (Ferrer-Admetlla et al., 2014). This analysis was limited to WGS data due to higher marker resolution of genomic regions, and presence of known phenotype classes. The XP-nSL statistic quantifies evidence of selection from patterns of elevated identity by state among haplotypes in a genomic region within one population relative to a reference population. This approach is robust to both recombination rate and demography and shows the potential to pick up soft sweeps and polygenic adaptation from standing variation (Ferrer-Admetlla et al., 2014; Szpiech et al., 2021). We calculated XP-nSL from phased and imputed vcf files from WGS samples between sea age classes for each regional grouping and sex, with 1SW individuals used as the reference grouping, using *selscan* 1.3.0 (Szpiech & Hernandez 2014), and normalized XP-nSL scores in 100Kb windows *norm* 1.3.0. Significant XP-nSL outliers were selected as those windows exceeding the 1% quantile for the greatest proportion of xpnSL scores greater than 2, indicating selective sweeps in multi-sea winter fish, or less than 2, indicating sweeps in 1SW fish. To estimate the extent that parallelism of genes with detected soft sweeps between groups occurred non-randomly, we carried out 10,000 random re-samples of the number of Entrez gene symbols from the *Salmo salar* gene list that were detected in the top 1% of comparisons for two compared groups (e.g., female fish in southern rivers, and female fish in northern rivers, and compared the observed rate of overlap against the randomly re-sampled distribution.

### Machine learning prediction of sea age from genotype dosage

As many genomic regions that fall below genome-wide statistical significance may still carry substantial predictive information (McGaugh et al., 2021), we next tested the utility of a machine learning approach to predict individual sea age using WGS data. We first selected the top 100, 200, 300, 400, and 500 loci with the highest likelihood ratio test score from each population-structure corrected GWA using WGS data to use as predictors in a random forest model (Breiman 2001), as in Hess et al., (2016) and suggested in Brieuc et al. (2018) to minimize overfitting. We then used continuous genotype dosages, which allow for quantifying the uncertainty at individual loci from differences in read depth or SNP quality for each locus (Zheng et al., 2011), as predictors of trait variation. Prediction of sea age for each sex and regional grouping was carried out using a random forest classification model (Breiman 2001) in the *randomForest* R package (Liaw & Weiner 2002), setting 25,000 trees and 80% of SNP sample size for the *mtry* parameter for each analysis. To estimate the prediction accuracy of each model, we estimated out-of-bag error (OOB), which quantifies the number of misidentified samples from a testing dataset with known categories. For each tree we specified a holdout of half of individuals from each sea age class, equalized by sample size, for testing and training, similar to the approach used by Briuec et al., (2018). We then compared the accuracy of the selected panel of SNPs by generating panels of 100-500 randomly selected SNPs, and calculated OOB for these panels to quantify classification error of random SNPs relative to those with elevated GWA significance. The trade-off between sensitivity and specificity was estimated for the full model of all samples for 500 SNP-prediction with a receiver operator characteristic (ROC) curve using the R package *pROC*.

### Gene set enrichment

To identify signals of polygenic selection at the pathway level, we next used the gene set enrichment approach implemented in the *polysel* set of R tools (Daub et al., 2013), which tests for shared patterns of elevated values of a population genetic parameter among genes within pathways. We downloaded gene information for Atlantic Salmon from **http://www.ncbi.nlm.nih.gov/gene**, and KEGG pathway information (Ogata et al., 1998) for Atlantic Salmon from https://www.ncbi.nlm.nih.gov/biosystems/. We then selected the maximum value of each tested statistic for each gene, and ran gene set enrichment on: Likelihood ratio test scores from genome-wide association with sea age in the WGS datasets, per-SNP PC1 scores from PCAngsd, -log10(*p*) *LFMM* scores from SNP array association with grilse proportion at K =1, and PC1 scores from *pcadapt*. We set a minimum of 5 genes per set, resulting in 237 sets and 11,437 genes analyzed for whole genome data, and 209 sets and 5507 genes analyzed for Axiom Array data. Significantly enriched KEGG pathways were identified using a *q* threshold of 0.05.

## Results

### Population structure

Population structure analyses using the SNP array uncovered structuring associated with regional differences in North American Atlantic Salmon populations (or 1SW salmon) (Figure 1A, B). Multiple genetic clusters below the regional level were also identified in PCA. We found that the first two PCs separated populations partially by regional differences in grilse proportion among sampled sites (primarily along the first PC axis, Supplementary Figure 1), explaining most of the genetic variation, after which percent variation explained fell below 2% (Supplementary Figure 2). Admixture analysis using *snmf* also supported multiple genetic clusters within broader geographic regions (Figure 1B, C).

We did not find genome-wide population structure associated with individual age classes of WGS individuals, in contrast to population-genetic clustering by river-level grilse proportion observed with the SNP array. Population structure inferred using genotype likelihoods in *PCAngsd* among the eight rivers in the whole genome re-sequencing dataset revealed regional separation between southern and northern rivers, and further clustering at the river level (Figure 2B). *PCAngsd* identified K=6 as optimally describing population structure (Figure 2C) and revealed more similar ancestry among close rivers within the same region.

### PCA-based signatures of selection

Using PCA-based scans for selection associated with population structure among populations genotyped on the SNP array, we uncovered many genomic regions exhibiting elevated divergence, including a previous sea-age associated locus, *six6* (Figure3A). We identified 3593 SNPs with elevated significance in PCA using *pcadapt* (*q* < 0.05), distributed genome-wide. These outliers overlapped 1139 genes (Supplementary Table 3), as well as the introgressed European karyotypic variant regions (Lehnert et al., 2018) on chromosomes ssa01and ssa23 (*n* = 559).

Selection scans of WGS samples using *PCAngsd* identified 52,986 significant SNPs overlapping 3262 genes (Supplementary Table 4), again including *six6* (Figure 4A). Significant association of population structure with karyotype variants was negligible (n=18, ssa01, n=1 ssa23) among WGS individuals.

### Genome-wide association with sea age

Using Axiom Array data and grilse proportions from 26 rivers across the North America, we found 11 associated SNPs at K=1, and one overlapping gene: *six6* (Figure 3B, Supplementary Table 5). This pattern of association at six6 was also strongest among the 976 SNPs, overlapping 312 distinct genes, found in the top 1% of RDA scores (Supplementary Table 6), and we found complete overlap of the 11 detected SNPs with K=1 and RDA and *pcadapt* outliers. However, after controlling for population structure in *LFMM* using K=5, we did not detect an association with *six6*, and instead identified six significantly associated SNPs overlapping 2 genes, with the strongest signal of association found at a tissue factor pathway inhibitor-like gene (Supplementary Table 5). We found both sets of SNPs detected with *LFMM* exhibited complete overlap with RDA outliers, and high overlap with and *pcadapt* (K1 = 11, 100%, K5 = 3, 50%) outliers. Variance partitioning of grilse proportion by principal component scores of genetic variation identified a significant (*p* < 0.001) and large proportion of variation explained by PC1 (60%), with smaller but significant contributions from PC2 (1%) and PC5 (4%).

**Figure 3.**
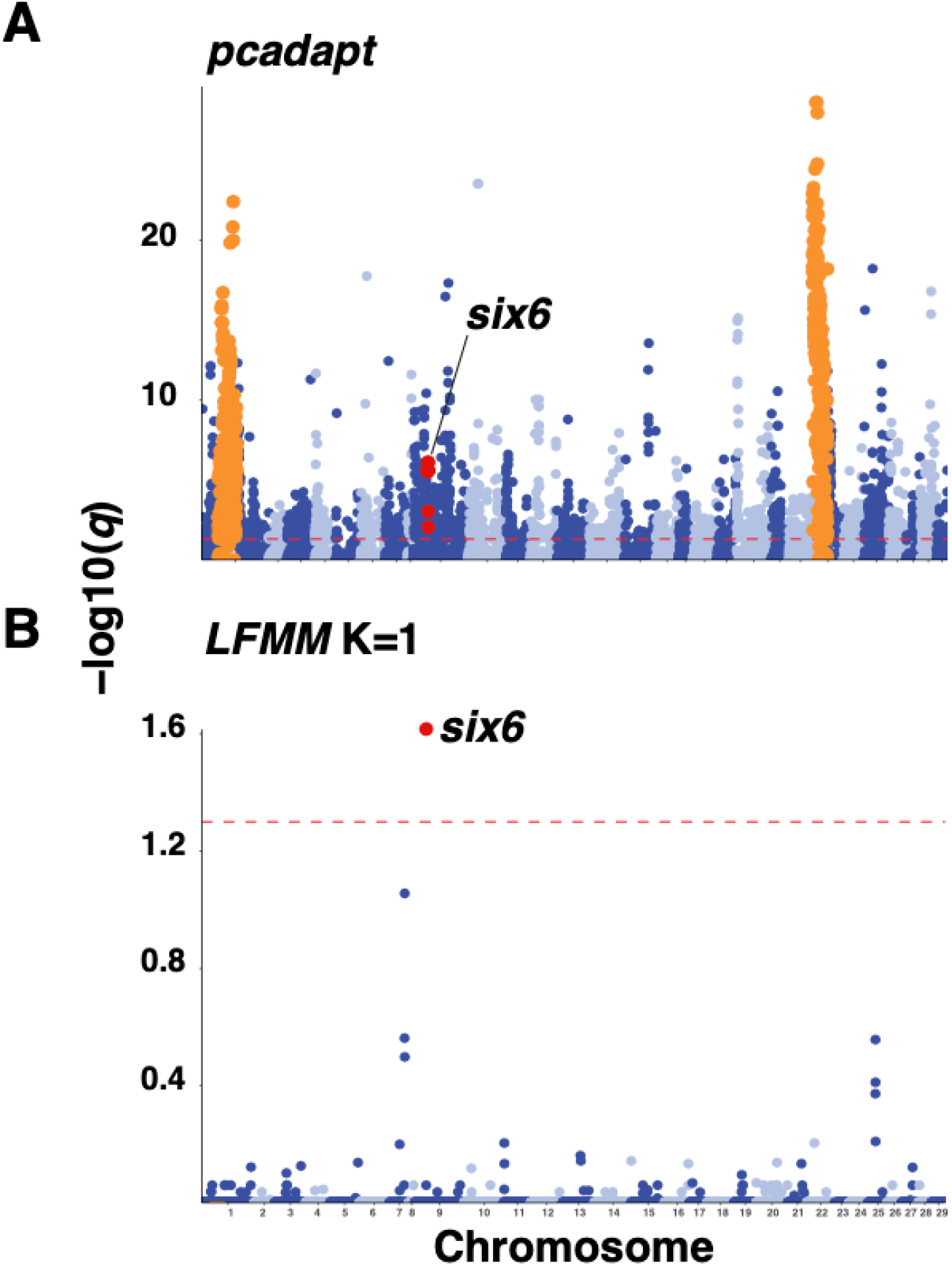
Genome-wide distribution of Axiom Array SNPs exhibiting significant association (red dotted line, *q* < 0.05) with population structure in *pcadapt* (A), grilse proportion in river of origin in LFMM with K = 1 (B) with red circles indicating the location of SNPs within *six6*, and orange circles indicating location of karyotypic variants identified by Lehnert et al. (2019)

Genome-wide association using individual phenotypes using the WGS data instead identified 32 significant loci, and an overlap with *vgll3* as the only significantly associated gene in the total dataset analysis (Figure 4B, Supplementary Table 7). In a finer-scale analysis of female fish from southern rivers, we identified 35 significant SNPs, overlapping with 3 genes: a predicted glutamate receptor, NMDA 2B-like on ssa02, a butyrophilin subfamily 2 member A1-like gene on ssa18, and *vgll3* (Figure 4C, Supplementary Table 6). In both comparisons, we uncovered a significant association at a genomic region on ssa25 showing the strongest sea age association in European Atlantic salmon, recently identified by Sinclair-Waters et al., (2021). No significant associations were identified within the other subsets at *q* < 0.05. In comparison with the genome-wide average, we identified elevated effect size (**β**) and *F*_ST_ values among the top 100 most significant SNPs, but no increased PC1 association (Table 1). Variance partitioning of sea age variation by PCs revealed a small but significant proportion of variance in individual sea age explained by PC1 (1.7%, *p* < 0.001), indicating reduced influence of population structure on individual level GWA statistics compared to SNP array data. Comparing *F*_ST_ to uncorrected GWA LRT revealed high and significant correlation (r^2^ = 0.914, *p* < 1 ×10^−15^), indicating imputation and genotype-likelihood estimates captured similar allele frequency variation across the genome

**Figure 4.**
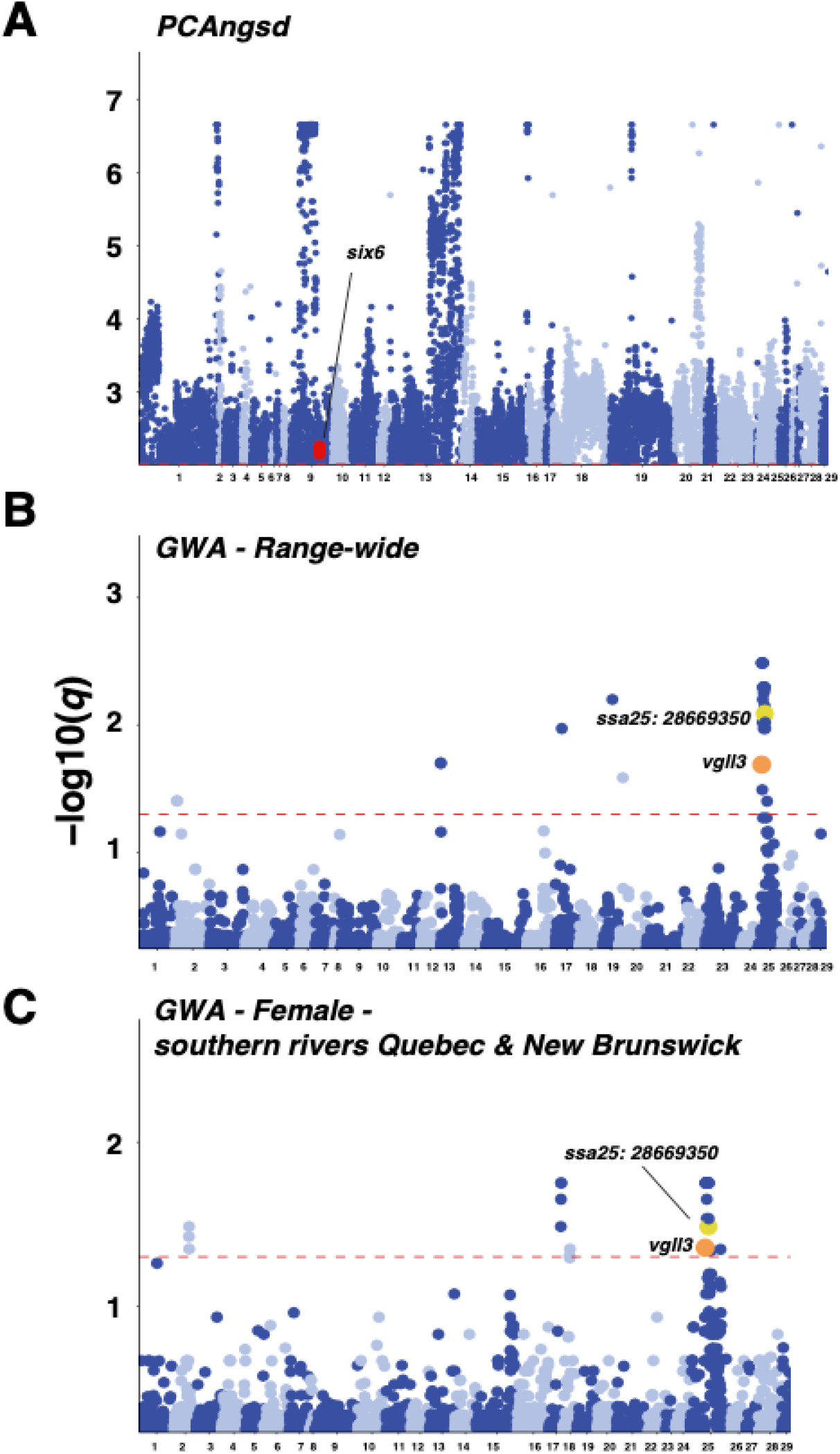
Genome-wide distribution of SNPs identified from low coverage whole genome re-sequencings exhibiting significant association (red dotted line, *q* < 0.05) with population structure in *PCAngsd* (A), individual sea age across all samples (B) individual sea age within female fish sampled in southern rivers in Quebec and New Brunswick Rivers (C), with red circles indicating the location of SNPs within *six6*, orange indicating within *vgll3*, and yellow indicating the location of the SNP with highest explanatory power for sea age on *ssa25* identified by Sinclair-Waters et al. (2021).

**Table 1.** *F*_ST_ between 1SW and MSW, GWA effect size (β)and PC1 loading in PCANGSD for the top 100 associated SNPs and genome-wide loci across separate GWA of sea age, with * denoting a significant difference in parameters between top SNP and genome-wide SNP comparisons.

| **GWA group** | **$F_{ST}$\*** |  | **Effect size ($\beta$)\*** |  | **PC1 loading** |  |
| --- | --- | --- | --- | --- | --- | --- |
|  | Top 100 SNPs | Genome-wide SNPs | Top 100 | Genome-wide SNPs | Top 100 SNPs | Genome-wide SNPs |
| Female – southern rivers Quebec and New Brunswick | 0.161 | 0.0007 | 3.19 | 0.38 | 0.759 | 1.02 |
| Male – southern rivers Quebec and New Brunswick | 0.0928 | 0.0002 | 9.05 | 0.335 | 0.945 | 1.02 |
| Female – northern rivers Quebec and Labrador | 0.0854 | 0.00009 | 4.27 | 0.311 | 0.923 | 1.02 |
| Male – northern rivers Quebec and Labrador | 0.1533 | 0.0003 | 4.40 | 0.418 | 0.677 | 1.04 |

### Detection of selection sweeps

Estimating XP-nSL to carry out haplotype-based selection scans, we identified a genome-wide distribution of genes in outlier windows in each comparison (Figure 5A, B, n=927-1000, Supplementary Table 8). We uncovered an association with a window containing *six6* distinct to northern rivers in both male and female fish (Figure 5A), and with *vgll3* in comparisons of female individuals from southern rivers, as well as *akap11*, another gene with previously identified associations with sea age (Barson et al., 2015, Figure 5B). Comparison of observed rates of overlap of top 1% of genes were low but exceeded random expectations (> 99.9^th^ percentile) in all comparisons. We identified the highest overlap in southern rivers (5.59%), and in females (3.34%) compared to in northern rivers (2.26%) and in male fish (2.02%).

**Figure 5.**
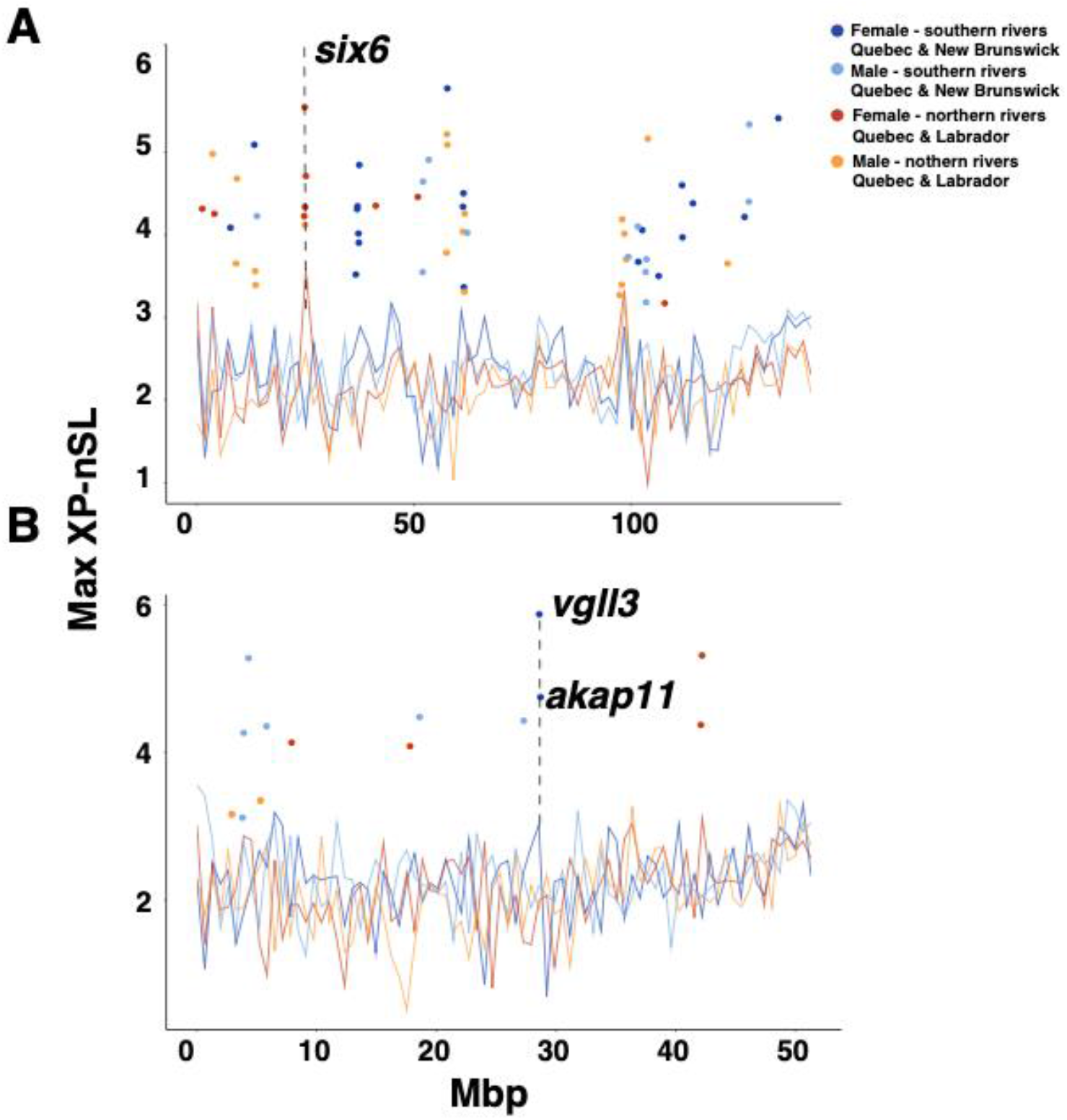
Smoothed XP-nSL scores and outliers in 100Kb windows for each sex and region at *ssa09* (A), and *ssa25*(B).

### Machine learning prediction of sea age

Across datasets, the 100 most significant SNPs based on LRT score could predict sea age with greater than 75% accuracy (Figure 6). Increasing the number of SNPs within each dataset increased accuracy to greater than 80% overall, and 90% in all but the Female southern rivers dataset and total dataset models. Investigation of the ROC curve for the total dataset identified both high sensitivity and specificity of random forest models using 500 sea age associated SNPs, inferred from rapid rise above the diagonal slope of a random classifier (Supplementary Figure 3). Across models with randomly selected SNPs, we found consistently low prediction accuracy (∼45%), even with the 500 SNP panel, indicating reduced predictive capacity of comparably sized panels of randomly selected genome-wide SNPs.

**Figure 6.**
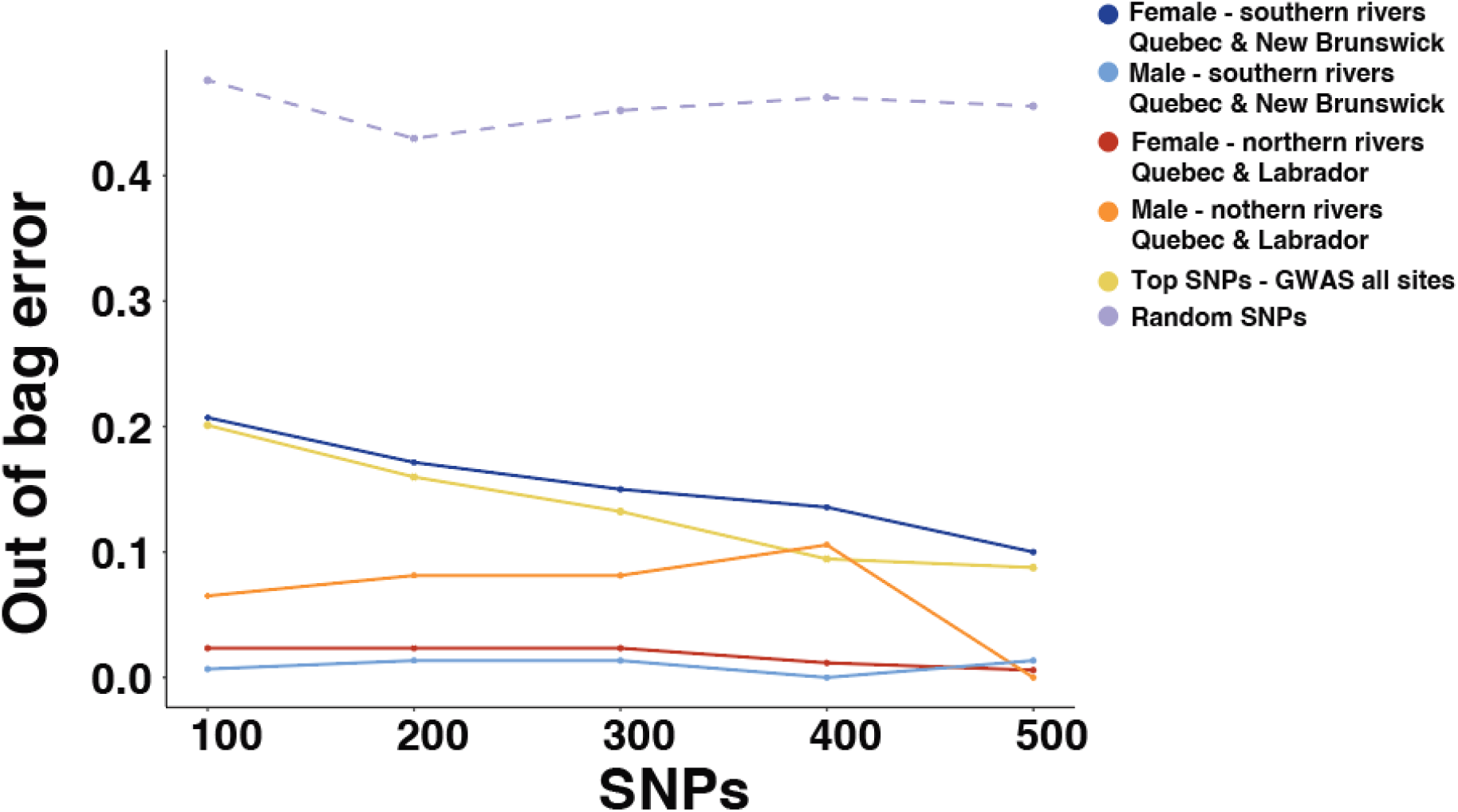
Random forest out of bag classification error rates across SNP panel sizes (100 – 500) for the highest ranking SNPS in GWAS for all samples stratified by sex and region. Randomly selected SNPs are represented with a dashed line.

### Gene set and gene ontology enrichment

Using gene set enrichment, we identified 37 significantly enriched KEGG pathways among GWA comparisons of individual sea age data, many of which were significantly enriched among multiple comparisons (Figure 7, Supplementary Table 9). In contrast, we did not identify significant enrichment among PC scores from WGS or SNP array analyses, or -log10(*p)* values from range-wide *LFMM* analyses. Hierarchical clustering of -log10(*q)* values from gene set enrichment revealed a core set of nine processes corresponding to developmental, osmoregulatory, neurological, and cardiovascular functions that were most significantly enriched among all groups, such as the phosphatidylinositol signalling system, adrenergic signalling in cardiomyocytes, the GnRH signaling pathway, and focal adhesion, We found the most significant enrichment among all groups in the calcium signalling pathway (Figure 7).

**Figure 7.**
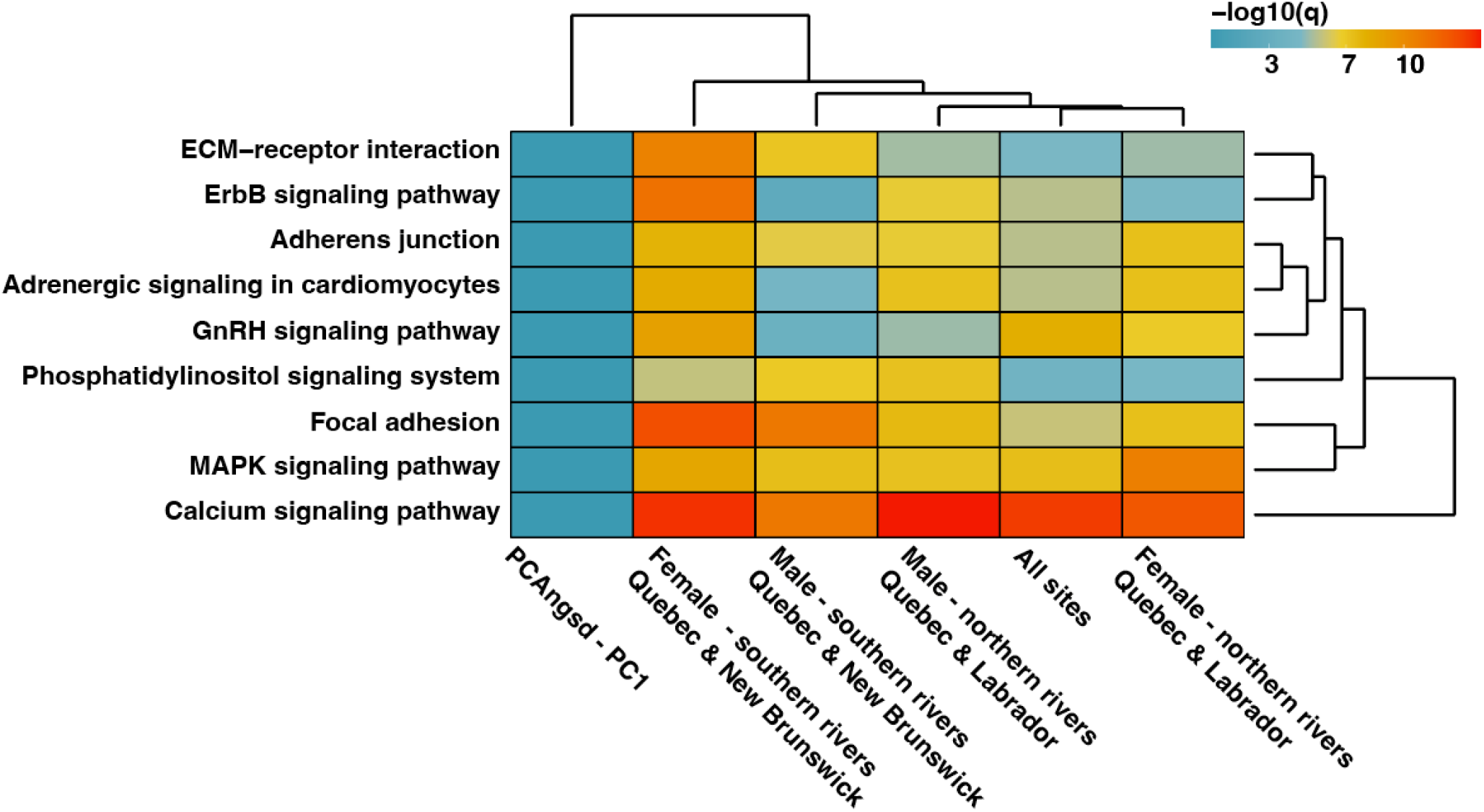
Hierarchical clustering of highly significant (*q* < 0.001) KEGG pathways identified using gene set enrichment across groups.

## Discussion

Identifying the genomic basis of adaptive traits in wild populations is key to understanding how evolutionary processes give rise to diverse and ecologically important phenotypes (Bolnick et al., 2018). Here, we characterized the genetic architecture of sea age variation in North American Atlantic Salmon using SNP array and WGS genomic datasets to test whether this genetic architecture is repeatable at two spatial scales: between European and North American populations, and within North American populations. Consistent with the expectation that large-effect loci will exhibit higher rates of parallelism (Barghi et al. 2020; Yeaman 2015), significant associations with sea age in North America overlapped with European sea age loci at the gene and SNP level (*vgll3, six6*). Within North America, we also found polygenic adaptation with low overlap of outliers, whereas gene set enrichment revealed repeatable patterns of association with conserved pathways associated with maturation. Our findings indicate contingency in the genetic architecture of sea age in Atlantic Salmon, but high parallelism of core molecular pathways and large-effect loci in generating the diversity of at-sea maturation strategies in Atlantic Salmon.

### Detection of known maturation loci

Our results across association analyses revealed parallel genetic architecture of sea age variation between North American and European Atlantic Salmon at the previously identified large-effect locus *vgll3*. Significant GWA association using WGS data paired with individual phenotypes, elevated scores for estimates of effect size, differentiation (*F*_ST_), and selective sweeps (XP-nSL) indicate strong selection at *vgll3* in female fish in southern rivers in Quebec and New Brunswick. This finding is concordant with past research in European populations, both in identifying strong selection and sea-age association at *vgll3*, and in finding sex-specific patterns of association and dominance at *vgll3* (Barson et al., 2015). Similar patterns of association with large-effect loci underlying sex-specific maturation and migration have also been revealed across the broader salmonid lineage, indicating large-effect loci may be common in genetic architectures for distinct life history strategies among salmonids (Pearse et al., 2019; Thompson et al., 2020).

Additionally, we uncovered a SNP-level pattern of parallelism at a *vgll3*-adjacent region between European and North American Atlantic Salmon. This region has previously been identified as highly significant in explaining sea age variation in European Atlantic Salmon in a recent candidate gene study (Sinclair-Waters et al., 2021). Our observation is concordant with the expectation that large-effect loci are more likely to exhibit parallelism (Yeaman 2015). The source of this parallelism, either through ancestral variation, repeated evolution, or migration, remains unknown, and is a clear goal for follow-up studies (Lee & Coop 2017). However, our results suggest that associations at *vgll3* and the *vgll3*-adjacent SNP varied across rivers and sexes, indicating heterogeneity in the genetic architecture of maturation across the North American range despite gene and SNP-level genetic parallelism. This finding is consistent with a past amplicon-based study (Kusche et al., 2017), which similarly identified significant variation in associations of *vgll3* and sea age dependent on sampled river. Variation in associations with *vgll3* and sea-age across North American rivers observed by Kusche et al. 2017 was attributed to a potential low proportion of MSW fish with advanced sea age (> 3SW) in amplicon-genotyped samples. Similarly, our sampling of fish across eight North American locations identified predominantly 2SW fish, and the small subset of fish with age greater than 2SW were repeat spawners with a 1-year virgin sea age (n = 11, 1.9%). This finding is concordant with the low proportion of older MSW fish among several rivers studied here (Kusche et al. 2017) and does not preclude *vgll3* exhibiting consistent associations with advanced sea age among rivers in North America as observed in European populations. Surprisingly, we do not find evidence of allelic associations at *vgll3* in population-level tests of association with grilse proportion, despite it being identified with individual re-sequencing data. The lack of this association at the population level may be a result of the sex and region-specific nature of this association, as well as the lack of known sea age classes in samples genotyped on the SNP array.

Using SNP array data, we found an association with river-level sea age and *six6*, which co-varied with population structure. The significant signal associated with population structure revealed in PCA, LFMM, and RDA analyses is also consistent with past associations of this locus with fine-scale population structure and river-specific features such as catchment area. This finding supports the possibility that this association may also correspond to a relationship with a correlated trait that is influenced by local environment rather than maturation itself (Pritchard et al., 2018; Zueva et al., 2021). This association may also reflect an artefact of sampling across regions exhibiting both population structure and differences in grilse proportion. GWA of individual level phenotypes and WGS data also failed to replicate this association in the present study, indicating *six6* itself may not be a major determinant of maturation across North American populations sampled here. Our identification of XP-nSL scores elevated in Northern male and female fish may indicate weak polygenic adaptation involving this locus in North American rivers, or selection acting on other targets within this 100KBp window. In contrast, GWA of a European multi-generation aquaculture line uncovered significant associations in genomic regions near *six6* (Sinclair-Waters et al., 2020), and this locus was also identified using haplotype-based and single SNP association models in follow-up analyses of wild individuals in Europe (Sinclair-Waters et al., 2021), indicating a more direct role for *six6* in maturation among European populations compared to North America.

Our combined results indicate high context-dependence of observed parallelism at *vgll3* and *six6*. These genes have also shown associations with maturation timing and growth across the broader vertebrate lineage, indicating a conserved genetic role in control of maturation (Cousminer et al., 2013; Perry et al., 2014). However, within salmonids, variable patterns of association have also been identified, with recurring associations with maturation across the lineage identified only at *six6* (Waters et al., 2021). We demonstrate that this relationship is variable even within Atlantic Salmon based on trans-Atlantic divergence, sex, and regional population structure. Similar patterns of variable reuse of key genes implicated in intraspecific ecological divergence have also been observed in studies of *Pungitius* and *Gasterosteus* sticklebacks (Fang et al., 2021,), as well as between *Timema* stick insect species (Villoutreix et al., 2020). Both variability in environmental conditions and drift likely plays a role in the reuse or detectability of *vgll3* and *six6*, as maturation has been shown to be impacted by population-specific genetic and environmental factors (Åsheim et al. 2022; Good & Davidson 2016; Mobley et al., 2021), and Atlantic Salmon populations exhibit both high drift and signals of environment-driven structuring (Bradbury et al., 2014). Together, these factors may generate variability in the underlying molecular mechanisms that generate individual maturation phenotypes and may not be shared between individuals or populations.

### Signals of polygenic adaptation

In addition to previously identified European large-effect loci, we uncovered genomic evidence of polygenic architecture underlying sea age variation across North American rivers. We build on recent detections of polygenic trait architecture underlying maturation detected within aquaculture (Mohamed et al., 2019; Sinclair-Waters et al., 2020) and breeding experiments (Debes et al., 2021) by identifying sets of predictive loci for maturation. We achieved high prediction accuracy when accounting for population and sex-level variation in per locus association with maturation and found non-parallel sets of loci enable accurate sea age prediction within regional groups. Consistent with other recent polygenic prediction studies in wild populations (Fuller et al., 2020; Hess et al., 2016; Lehnert et al., 2019), our results highlight the utility of prediction approaches when accounting for genotype uncertainty, and reveal a high capacity to predict sea age in wild Atlantic Salmon populations from genomic data.

Cross-population scans of soft sweeps uncovered candidate loci from previous studies and GWA comparisons (*vgll3, six6, akap11*), and low but significant overlap (2-5%) of genes among the most divergent haplotypes, consistent with population-specific polygenic adaptation. We found a largely non-parallel, but also non-random genomic basis of maturation at small effect loci across North American Atlantic Salmon populations. Similar to our observation of parallelism at large-effect loci, these findings suggest a role for local adaptation and contingency at the genomic level in the evolution of life history diversity. Given the diversity of habitats colonized by Atlantic Salmon (Klemetsen et al., 2003), local environmental variation could play a role in biasing parallelism of polygenic trait architectures and has been shown to significantly drive genome-wide population structuring in Atlantic Salmon (Bradbury et al., 2014; Moore et al., 2014). Population structure itself has also been shown to constrain rates of parallelism and promote heterogeneity in genetic architecture through stochastic loss of standing variation (Fang et al., 2021), and identification of hierarchical population structure and high post-colonization drift in this and past studies is consistent with this process (Bradbury et al., 2014). Lastly, genome duplications may facilitate non-parallelism through diversification from duplicated gene copies (Ohno 1970), and whole genome duplication within salmonids (Allendorf & Thorgard 1984; Lien et al., 2016) may have provided many genomic substrates for adaptation (Campbell et al., 2021)).

### Enrichment of shared processes

Our findings from gene set enrichment tests (Daub et al., 2013) indicate significant enrichment in pathways and molecular processes associated with maturation across the broader vertebrate lineage. Parallel enrichment at calcium signalling pathways is consistent with a significant role for calcium metabolism during oocyte maturation (Tosti 2006), and osmoregulation changes during river spawning (Persson et al., 1998). Additional pathways also revealed significant associations with sexual maturation, including MAPK signalling, which plays a role in meiotic maturation (Kishimoto 2003), Phosphatidylinositol signalling associated with oocyte maturation, and previously in association with polygenic sea age variation in aquaculture fish (Hoshino et al., 2004, Mohamed et al., 2019), and gonadotropin releasing hormone (GnRH), implicated in male maturation (Wen et al., 2010). Neurological processes which may underlie necessary behavioural changes for both sexual maturation and migration (Mobley et al., 2021) were also enriched across comparisons, and we find enrichment in ErbB signalling, which has been associated with neural development and aggression (Barros et al., 2009), and Phosphatidylinositol signalling, which has also been shown to mediate nervous system function (Raghu et al., 2019). Detection of significant enrichment among pathways implicated in cardiovascular function are also consistent with differences in energetic costs of large body size and migration in older sea winter fish (Jonsson et al 1997), and elevated *vgll3* expression in heart tissue during maturation (Verta et al., 2020).

We find a high degree of parallelism at the pathway level despite low gene reuse across North American populations, consistent with increasing parallelism at higher levels of organization. Studies of parallelism across traits have frequently identified high parallelism at the level of integrated traits, that declines at finer scales of comparison (Bolnick et al., 2018). At the molecular level, recent investigations into parallelism in other salmonids (Jacobs et al., 2020) and humans (Bergey et al., 2018) have similarly uncovered a high degree of parallelism of enriched pathways despite low genetic overlap, suggesting that polygenic architectures underlying complex traits may map on to relatively limited molecular processes. Our results indicate largely idiosyncratic genomic changes at the gene and SNP level across regions and sexes contribute to a few shared higher-level molecular pathways, corresponding to discrete differences in sea age. The observed molecular parallelism at the pathway level indicates that conserved processes with associations in other vertebrate lineages also underlie maturation in Atlantic Salmon.

### Limitations and future directions

Our study provides the first genomic investigation into polygenic and molecular components of sea age variation in North American Atlantic Salmon, but follow-up studies are necessary to build on uncertainty from limitations to the approaches used here. Our identification of different genes associated with sea age variation across individual level (WGS) and site-level (SNP array) datasets indicate remaining sensitivity to population structure in analyses of population-level phenotypic information. Replicability and informativeness of GWA studies can be significantly reduced when population structure is not appropriately controlled (Korte & Farlow 2013); this challenge is exacerbated in instances of overlap between population structure and adaptation making population structure correction overly conservative in some instances (Francois et al., 2016). However, our individual level re-sequencing showed only a weak association with major axes of population structure, and across methods we uncovered previously identified large-effect sea age loci, highlighting the utility of range-wide sampling and individual phenotypes for GWA in wild populations. However, our sample sizes here are smaller than large scale human and agricultural GWA studies, and follow-up validation with larger, independent datasets and meta-analysis of GWA scores will provide greater certainty.

An additional potential source of uncertainty is that our WGS analyses were restricted to low depth (∼4x) resequenced SNPs. Recent studies indicate significant roles for structural and epigenomic variation in mediating adaptation across many systems (Layton & Bradbury 2021), and future assay of these types of variation may identify additional molecular mechanisms important in sea age variation. Low depth re-sequencing also introduces some uncertainty about variants detected. However, we employed large sample size (n > 500), and statistical control for genotype uncertainty through genotype likelihood approaches and imputation. Recent simulations have shown that with sample sizes, sequencing depths, and levels of population structure used here, both likelihood and imputation approaches should accurately capture allele frequencies (Lou et al., 2021), and we find very high and significant correlation among *F*_ST_ from phased data and genotype-likelihood GWA scores. However, as sequencing quality and depth improves, sequencing larger panels of individuals with technologies that capture more classes of variant (e.g. epigenomic modifications, structural variants) will aid in characterizing the genomic basis of sea age variation.

## Conclusion

Adaptation of complex phenotypes such as life history variation often present comparably complex underlying architectures. These traits may evolve through a combination of evolution from standing variation, hard and soft sweeps, and alleles of varying effect size. Here, we used GWA, genome scans, sweep detection, and random forest-based polygenic prediction to explore the genomic basis of sea age variation in North American Atlantic Salmon. We found evidence of significant association with previously identified large-effect genes (*vgll3, six6*), as well as varying patterns of association based on both sex and region, suggesting both parallelism at large-effect loci, and a high degree of genetic redundancy and polygenicity in this trait. Despite low overlap of the most strongly associated genes, we found a set of molecular pathways with conserved roles in maturation were consistently enriched among comparisons, revealing a core set of molecular mechanisms that underlie sea age variation in Atlantic Salmon. Our findings demonstrate clear pathways and genes for future investigations of maturation traits, and show how methods aimed at resolving polygenic patterns can uncover the molecular basis of a complex phenotype in wild populations.

## Acknowledgements

We thank DFO staff and private partners for sampling and scale-reading for sea age inference, CIGENE for SNP genotyping and data processing, and Genome Quebec for whole genome sequencing. Thanks to Brendan Wringe for assistance with regional assignments of SNP array populations. This study was supported by the Genomics Research and Development Initiative (GRDI) and the Program for Aquaculture Regulatory Research (PARR) of the Department of Fisheries and Oceans Canada (DFO) and the Natural Sciences Engineering and Research Council Canada (NSERC). Samples for the Restigouche River were generously donated by Listuguj Mi’gmaq fishers to GMRC to contribute to the advancement of Atlantic salmon research.

## Author contributions

TK, SJD, AM, and IRB designed the study. Sampling and fieldwork were carried out by SJD and CAG. Laboratory work for sequencing was carried out by AM. Grilse proportions were estimated by JBD, GC, JA, CB, and SJL. TK conducted all statistical analyses of genomic data with assistance by CW, JN, CN, and SJL, and drafted the manuscript. All authors contributed to writing and editing of the manuscript.

### Data Availability

SNP array genotypes in plink format, and phenotype data for association analyses will be made available on dryad. Raw reads from WGS data will be uploaded to the NCBI SRA. All scripts used for analysis in this study are available at: https://github.com/TonyKess/seaage_GWAS

## References

Allendorf F. W. & Thorgaard, G. H. (1994). Tetraploidy and the evolution of salmonid fishes. In: Evolutionary genetics of fishes pp. 1–53. Springer, Boston, MA.

Andrews, S. (2010). FastQC: a quality control tool for high throughput sequence data. Available online at: http://www.bioinformatics.babraham.ac.uk/projects/fastqc

Åsheim, E.R., Debes, P.V., House, A., Niemelä, P.T., Siren, J.P., Erkinaro, J., Primmer, C.R. (2022) Strong effects of temperature, population and age-at-maturity genotype on maturation probability for Atlantic salmon in a common garden setting. bioRxiv.doi: https://doi.org/10.1101/2022.07.22.501167

Barghi, N., Tobler, R., Nolte, V., Jakšić, A. M., Mallard, F., Otte, K. A., Dolezal, M., Taus, T., Kofler, R., & Schlötterer, C. (2019). Genetic redundancy fuels polygenic adaptation in Drosophila. PLoS Biology, 17(2), e3000128.

Barros, C. S., Calabrese, B., et l. (2009). Impaired maturation of dendritic spines without disorganization of cortical cell layers in mice lacking NRG1/ErbB signaling in the central nervous system. Proceedings of the National Academy of Sciences, 106(11), 4507–4512.

Barson, N. J., Aykanat, T., Hindar, K. Baranski, et al. (2015) Sex-dependent dominance at a single locus maintains variation in age at maturity in salmon. Nature, 528(7582):405–408.

Bergey, C. M., Lopez, M., Harrison, G. F., Patin, E., Cohen, J. A., Quintana-Murci, L., Barreiro, L. B. & Perry, G. H. (2018). Polygenic adaptation and convergent evolution on growth and cardiac genetic pathways in African and Asian rainforest hunter-gatherers. Proceedings of the National Academy of Sciences, 115(48). 11256–11263.

Blount, Z.D., Lenski, R. E., & Losos, J. B. (2018). Contingency and determinism in evolution: Replaying life’s tape. Science, 362, eaam5979.

Bolnick, D. I., Barrett, R. D., Oke, K. B., Rennison, D. J., & Stuart, Y.E. (2018). (Non) parallel evolution. Annual Review of Ecology, Evolution, and Systematics, 49, 303–330.

Boulding, E. G., Ang, K. P., Elliott, J. A., Powell, F. & Schaeffer, L. R. (2019) Differences in genetic architecture between continents at a major locus previously associated with sea age at sexual maturity in European Atlantic salmon. Aquaculture, 500, 670–678.

Bradbury, I. R, Hamilton, L. C, Robertson, M. J., Bourgeois, C. E., Mansour, A. & Dempson, J. B. (2014). Landscape structure and climatic variation determine Atlantic salmon genetic connectivity in the Northwest Atlantic. Canadian Journal of Fisheries and Aquatic Sciences, 71(2), 246–258.

Breiman, L. (2001). Random forests. Machine Learning. 45(1), 5–32.

Brieuc, M. S., Waters, C. D., Drinan, D. P. & Naish, K. A. (2018). A practical introduction to Random Forest for genetic association studies in ecology and evolution. Molecular Ecology Resources, 18(4), 755–766.

Browning, S. R. & Browning, B. L. (2007) Rapid and accurate haplotype phasing and missing data inference for whole genome association studies by use of localized haplotype clustering. American Journal of Human Genetics, 81, 1084–1097.

Campbell, M. A., Anderson, E. C., Garza, J. C. & Pearse, D. E.. (2021) Polygenic basis and the role of genome duplication in adaptation to similar selective environments. Journal of Heredity, 112(7), 614–625.

Caye, K., Jumentier, B., Lepeule, J. & François, O. (2019). LFMM 2: fast and accurate inference of gene-environment associations in genome-wide studies. Molecular Biology and Evolution, 36(4), 852–860.

Conte, G. L., Arnegard, M. E, Peichel, C. L., & Schluter, D. (2012). The probability of genetic parallelism and convergence in natural populations. Proceedings of the Royal Society B: Biological Sciences, 279(1749), 5039–5047.

Cousminer, D. L., Berry, D. J., Timpson, N. J., Ang, W., Thiering, E., Byrne, E. M., et al. (2013). Genome-wide association and longitudinal analyses reveal genetic loci linking pubertal height growth, pubertal timing and childhood adiposity. Human Molecular Genetics, 22, 2735–2747.

Czorlich, Y., Aykanat, T., Erkinaro, J., Orell, P., & Primmer, C. R. (2018). Rapid sex-specific evolution of age at maturity is shaped by genetic architecture in Atlantic salmon. Nature Ecology & Evolution, 2(11), 1800–1807.

Dadswell, M., Spares, A., Reader, J., McLean, M., McDermott, T., Samways, K., & Lilly, J. (2021). The decline and impending collapse of the Atlantic Salmon (Salmo salar) population in the North Atlantic Ocean: a review of possible causes. Reviews in Fisheries Science & Aquaculture. 30(2), 215 – 218.

Danecek, P., Auton, A., Abecasis, G., Albers, C.A., Banks, E., DePristo, M.A., Handsaker, R.E., Lunter, G., Marth, G.T., Sherry, S.T., McVean, G. (2011). The variant call format and VCFtools. Bioinformatics, 27, 2156–2158.

Daub, J.T., Hofer, T., Cutivet, E., Dupanloup, I., Quintana-Murci, L., Robinson-Rechavi, M., & Excoffier, L. (2013). Evidence for polygenic adaptation to pathogens in the human genome. Molecular Biology and Evolution, 30(7), 1544–1558.

Debes, P. V., Piavchenko, N., Ruokolainen, A., Ovaskainen, O., Moustakas-Verho, J. E., Parre, N., Aykanat, T., Erkinaro, J., & Primmer, C. R. (2021) Polygenic and major-locus contributions to sexual maturation timing in Atlantic salmon. Molecular Ecology, 30(18), 4505–4519.

DePristo, M.A., Banks, E., Poplin, R., Garimella, K.V., Maguire, J.R., Hartl, C., Philippakis, A.A., Del Angel, G., Rivas, M.A., Hanna, M., McKenna, A. (2011). A framework for variation discovery and genotyping using next-generation DNA sequencing data. Nature Genetics, 43(5), 491.

Elmer, K. R., & Meyer, A. (2011). Adaptation in the age of ecological genomics: Insights from parallelism and convergence. Trends in Ecology & Evolution, 26, 298–306.

Fagny, M. & Austerlitz, F. (2021). Polygenic adaptation: integrating population genetics and gene regulatory networks. Trends in Genetics, 7(7), 631–638.

Fang, B., Kemppainen, P., Momigliano, P. & Merilä, J. (2021). Population structure limits parallel evolution in sticklebacks. Molecular Biology and Evolution, 38(10), 4205–4221.

François, O., Martins, H., Caye, K., & Schoville, S. D. (2016). Controlling false discoveries in genome scans for selection. Molecular Ecology, 25(2), 454–469.

Frichot, E. & Francois, O. (2015). LEA: An R Package for landscape and ecological association studies. Methods in Ecology and Evolution, 6, 925–929.

Frichot, E., Mathieu, F., Trouillon, T., Bouchard, G., François, O. (2014). Fast and efficient estimation of individual ancestry coefficients. Genetics, 196, 973–983.

Friedland, K. D,, Hansenm L. P., Dunkley, D. A,, & MacLean, J. C. (2000). Linkage between ocean climate, post-smolt growth, and survival of Atlantic salmon (Salmo salar L.) in the North Sea area. ICES Journal of Marine Science, 57(2), 419–429.

Fuller, Z. L., Mocellin, V. J., Morris, L. A., Cantin, N., et al. (2020). Population genetics of the coral Acropora millepora: Toward genomic prediction of bleaching. Science, 369(6501):eaba4674.

Garant, D., Dodson, J. J., & Bernatchez, L. (2003). Differential reproductive success and heritability of alternative reproductive tactics in wild Atlantic salmon (Salmo salar L.). Evolution, 57(5), 1133–1141.

Good, C. & Davidson, J. (2016) A review of factors influencing maturation of Atlantic salmon, Salmo salar, with focus on water recirculation aquaculture system environments. Journal of the World Aquaculture Society, 47(5), 605–632.

Hess, J. E., Zendt, J. S., Matala, A. R., Narum, S. R. (2016) Genetic basis of adult migration timing in anadromous steelhead discovered through multivariate association testing. Proceedings of the Royal Society B: Biological Sciences, 283(1830), 20153064.

Hoban, S., Kelley, J. L., Lotterhos, K. E., Antolin, M. F., Bradburd, G., Lowry, D. B., Poss, M. L., Reed, L. K., Storfer, A., & Whitlock, M.C. (2016). Finding the genomic basis of local adaptation: pitfalls, practical solutions, and future directions. The American Naturalist, 188(4), 379–97.

Hoshino, Y., Yokoo, M., Yoshida, N., Sasada, H., Matsumoto, H. & Sato, E. (2004). Phosphatidylinositol 3-kinase and Akt participate in the FSH-induced meiotic maturation of mouse oocytes. Molecular Reproduction and Development: Incorporating Gamete Research, 69(1), 77–86.

Hutchings, J. A. & Jones, M. E. (1998). Life history variation and growth rate thresholds for maturity in Atlantic salmon, Salmo salar. Canadian Journal of Fisheries and Aquatic Sciences, 55(S1), 22–47.

Jacobs, A., Carruthers, M., Yurchenko, A., Gordeeva, N.V, Alekseyev, S., Hooker, O., Leong, J., Minkley, D.R., Rondeau, E., Koop, B., Adams, C., Elmer, K.R. (2020) Parallelism in eco-morphology and gene expression despite variable evolutionary and genomic backgrounds in a Holarctic fish. PLoS Genetics, 16(4): e1008658.

Jeffery, N. W., Stanley, R. R., Wringe, B. F., Guijarro-Sabaniel, J., Bourret, V., Bernatchez, L., Bentzen, P., Beiko, R. G., Gilbey, J., Clément, M., & Bradbury, I. R. (2017). Range-wide parallel climate-associated genomic clines in Atlantic salmon. Royal Society Open Science, 4(11), 171394.

Johnston, S. E., Orell, P., Pritchard, V. L., Kent, M.P., Lien, S., Niemelä, E., Erkinaro, J., & Primmer, C. R. (2014) Genome wide SNP analysis reveals a genetic basis for sea age variation in a wild population of Atlantic salmon (Salmo salar). Molecular Ecology, 23(14), 3452–3468.

Jonsson N., & Jonsson, B. (2007). Sea growth, smolt age and age at sexual maturation in Atlantic salmon. Journal of Fish Biology, 71(1), 245–252.

Jonsson, N., Jonsson, B., & Hansen, L. P. (1997). Changes in proximate composition and estimates of energetic costs during upstream migration and spawning in Atlantic salmon Salmo salar. Journal of Animal Ecology, 66, 425–436.

Jørsboe, E. & Albrechtsen, A. (2022) Efficient approaches for large-scale GWAS with genotype uncertainty, G3 Genes|Genomes|Genetics, 12(1), jkab385, https://doi.org/10.1093/g3journal/jkab385

Kaeuffer, R., Peichel, C. L., Bolnick, D. I., & Hendry, A. P. (2012). Parallel and nonparallel aspects of ecological, phenotypic, and genetic divergence across replicate population pairs of lake and stream stickleback. Evolution, 66(2), 402–418.

Kishimoto, T. (2003). Cell-cycle control during meiotic maturation. Current Opinion in Cell Biology, 15(6), 654–663.

Klemetsen, A., Amundsen, P. A., Dempson, J. B., Jonsson, B., Jonsson, N., O’connell, M. F., & Mortensen, E. (2003). Atlantic salmon Salmo salar (L)., brown trout Salmo trutta (L). and Arctic charr Salvelinus alpinus (L.): a review of aspects of their life histories. Ecology of Freshwater Fish, 12(1), 1–59.

Kreiner, J. M., Tranel, P. J., Weigel, D., Stinchcombe, J. R., & Wright, S. I. (2021). The genetic architecture and population genomic signatures of glyphosate resistance in Amaranthus tuberculatus. Molecular Ecology, 30(21), 5373–5389.

Korte A. & Farlow, A. (2013). The advantages and limitations of trait analysis with GWAS: a review. Plant Methods, 9(1), 1–9.

Kusche, H., Côté, G., Hernandez, C., Normandeau, E., Boivin Delisle, D., & Bernatchez, L. (2017). Characterization of natural variation in North American Atlantic Salmon populations (Salmonidae: Salmo salar) at a locus with a major effect on sea age. Ecology and Evolution, 7(15), 5797–807.

Layton, K. K., & Bradbury, I. R. (2021). Harnessing the power of multi-omics data for predicting climate change response. Journal of Animal Ecology.

Lee, K. M. & Coop, G. (2017). Distinguishing among modes of convergent adaptation using population genomic data. Genetics, 207(4), 1591–1619.

Lehnert, S.J., Kess, T., Bentzen, P., Kent, M.P., Lien, S., Gilbey, J., Clément, M., Jeffery, N.W., Waples, R.S. and Bradbury, I.R., 2019. Genomic signatures and correlates of widespread population declines in salmon. Nature Communications, 10(1), pp.1–10.

Lehnert, S. J., Kess, T., Bentzen, P., Clément, M., & Bradbury, I. R. (2020). Divergent and linked selection shape patterns of genomic differentiation between European and North American Atlantic salmon (Salmo salar). Molecular Ecology, 29(12), 2160–2175.

Lehnert, S. J., Bentzen, P., Kess, T., Lien, S., Horne, J. B., Clément, M., & Bradbury, I. R. (2019). Chromosome polymorphisms track trans Atlantic divergence and secondary contact in Atlantic salmon. Molecular Ecology, 28(8):2074–2087.

Li, H. (2013). Aligning sequence reads, clone sequences and assembly contigs with BWA-MEM. arXiv, 1303, 3997.

Li, H., Handsaker, B., Wysoker, A., Fennell, T., Ruan, J., Homer, N., Marth, G., Abecasis, G., Durbin, R. (2009). The sequence alignment/map format and SAMtools. Bioinformatics; 25(16), 2078–2079.

Liaw, A. & Wiener M. (2002). Classification and regression by randomForest. R News, 2(3), 18–22.

Lien, S., Koop, B. F., Sandve, S. R., Miller, J. R., Kent, M. P., Nome, T., et al. (2016). The Atlantic salmon genome provides insights into rediploidization. Nature, 533, 200–205.

Lou, R. N., Jacobs, A., Wilder, A. P., & Therkildsen, N. O. (2021). A beginner’s guide to low-coverage whole genome sequencing for population genomics. Molecular Ecology, 30(23), 5966–5993.

Martin, M. (2011) Cutadapt removes adapter sequences from high-throughput sequencing reads. EMBnet. Journal, 17(1),10–12.

Mayjonade, B., Gouzy, J., Donnadieu, C., Pouilly, N., Marande, W., Callot, C., Langlade, N., Muños, S. (2016). Extraction of high-molecular-weight genomic DNA for long-read sequencing of single molecules. BioTechniques 61(4), 203–205.

McGaugh, S. E., Lorenz, A. J. & Flagel, L.E. (2021). The utility of genomic prediction models in evolutionary genetics. Proceedings of the Royal Society B. 288(1956), 20210693.

Meisner, J. & Albrechtsen, A. (2018). Inferring population structure and admixture proportions in low-depth NGS data. Genetics, 210(2), 719–731.

Mobley, K.B., Aykanat, T., Czorlich, Y., House, A., Kurko, J., Miettinen, A., Moustakas-Verho, J., Salgado, A., Sinclair-Waters, M., Verta, J. P., & Primmer, C.R. (2021) Maturation in Atlantic salmon (Salmo salar, Salmonidae): a synthesis of ecological, genetic, and molecular processes. Reviews in Fish Biology and Fisheries, 31, 523 – 571.

Mohamed, A. R., Verbyla, K. L., Al-Mamun, H. A., McWilliam, S., Evans, B., King, H., Kube, P., & Kijas, J. W. (2019). Polygenic and sex specific architecture for two maturation traits in farmed Atlantic salmon. BMC Genomics, 20, 139, https://doi.org/10.1186/s12864-019-5525-4

Moore, J. S., Bourret, V., Dionne, M., Bradbury, I. R., O’Reilly, P., Kent, M., Chaput, G., & Bernatchez L. (2014) Conservation genomics of anadromous Atlantic salmon across its North American range: outlier loci identify the same patterns of population structure as neutral loci. Molecular Ecology, 23(23), 5680–5697.

Ohno, S. (1970). The enormous diversity in genome sizes of fish as a reflection of nature’s extensive experiments with gene duplication. Transactions of the American Fisheries Society, 99(1),120–130.

Ogata, H., Goto, S., Fujibuchi, W. & Kanehisa, M. (1998) Computation with the KEGG pathway database. Biosystems. 47(1-2), 119–28.

Oksanen, J., Guillaume, F. G., Friendly, M., Kindt, R., Legendre, P. McGlinn, D. Minchin, P. R., O’Hara, R. B., Simpson, G. L., Solymos, P., Stevens, M. H. H., Szoecs, E. & Wagner, H. (2020). vegan: Community Ecology Package. R package version 2.5-7. https://CRAN.R-project.org/package=vegan

Olmos, M., Payne, M.R., Nevoux. M/, Prévost. E/, Chaput. G/, Du Pontavice. H/, Guitton. J., Sheehan, T., Mills, K., & Rivot, E. (2020), Spatial synchrony in the response of a long range migratory species (Salmo salar) to climate change in the North Atlantic Ocean. Global Change Biology, 26, 1319–1337.

Pearse, D. E., Barson, N. J., Nome, T., Gao, G., Campbell, M. A., et al. (2019) Sex-dependent dominance maintains migration supergene in rainbow trout. Nature Ecology & Evolution, 3(12), 1731–1742.

Perry, J. R. B., Day, F., Elks, C. E., Sulem, P., Thompson, D. J., Ferreira, T., et al. (2014). Parent-of-origin-specific allelic associations among 106 genomic loci for age at menarche. Nature, 514: 92–97.

Ralph, P. L. & Coop, G. (2015). The role of standing variation in geographic convergent adaptation. The American Naturalist, 186(S1), S5–S23.

Persson, P., Sundell, K., Björnsson, B. T. & Lundqvist, H.. (1998). Calcium metabolism and osmoregulation during sexual maturation of river running Atlantic salmon. Journal of Fish Biology, 52(2), 334–349.

Pritchard, V. L., Mäkinen, H., Vähä, J. P., Erkinaro, J., Orell, P. & Primmer, C.R. (2018). Genomic signatures of fine-scale local selection in Atlantic salmon suggest involvement of sexual maturation, energy homeostasis and immune defence-related genes. Molecular Ecology, 27(11), 2560–75.

Privé, F., Luu, F., Vilhjálmsson, B. J., & Blum, M. G. B. (2020). Performing highly efficient genome scans for local adaptation with R package pcadapt version 4. Molecular Biology and Evolution, 37(7), 2153–2154.

Raghu, P., Joseph, A., Krishnan, H., Singh, P. & Saha, S. (2019. Phosphoinositides: regulators of nervous system function in health and disease. Frontiers in Molecular Neuroscience, 12, 208.

Rougemont, Q. & Bernatchez, L. (2018). The demographic history of Atlantic salmon (Salmo salar) across its distribution range reconstructed from approximate Bayesian computations. Evolution, 72(6), 1261–1277.

Salisbury, S., McCracken, G. R., Perry, R., Keefe, D., Layton, K. K., Kess, T., Nugent, C. M., Leong, J. S., Bradbury, I. R., Koop, B. F., Ferguson, M.M. & Ruzzante, D. E. (2022) The genomic consistency of the loss of anadromy in an Arctic fish (Salvelinus alpinus). The American Naturalist, 199(5).

Sinclair-Waters, M., ØdegÅrd, J., Korsvoll, S. A., Moen, T., Lien, S., Primmer, C. R., & Barson, N. J. (2020). Beyond large-effect loci: large-scale GWAS reveals a mixed large-effect and polygenic architecture for age at maturity of Atlantic salmon. Genetics Selection Evolution, 52, 9.

Sinclair-Waters, M., Nome, T., Wang, J., Lien, S., Kent, M. P., Sægrov, H., Florø-Larsen, B., Bolstad, G. H., Primmer, C. R. & Barson, N. J. (2021) Dissecting the loci underlying maturation timing in Atlantic salmon using haplotype and multi-SNP based association methods. bioRxiv.

Storey, J.D., Bass, A.J., Dabney, A., Robinson, D. (2015). qvalue: Q-value estimation for false discovery rate control. R package version 2.10.0.http://github.com/jdstorey/qvalue

Storey J.D., & Tibshirani, R. (2003) Statistical significance for genomewide studies. Proceedings of the National Academy of Sciences of the United States of America, 100, 9440–9445.

Szpiech, Z. A., Novak, T. E., Bailey, N. P., Stevison, L. S. (2021) Application of a novel haplotype-based scan for local adaptation to study high-altitude adaptation in rhesus macaques. Evolution Letters, 5(4), 408–421.

Szpiech ZA, & Hernandez RD. (2014). selscan: an efficient multithreaded program to perform EHH-based scans for positive selection. Molecular Biology and Evolution, 31(10), 2824–2827.

Therkildsen, N.O. & Palumbi, S.R. (2017). Practical low-coverage genome-wide sequencing of hundreds of individually barcoded samples for population and evolutionary genomics in non-model species. Molecular Ecology Resources, 17(2), 194–208.

Thompson, N. F., Anderson, E. C., Clemento, A. J., Campbell, M. A., Pearse, D. E., Hearsey, J. W., Kinziger, A. P., & Garza, J.C. (2020) A complex phenotype in salmon controlled by a simple change in migratory timing. Science, 370(6516), 609–613.

Tosti,E. (2006). Calcium ion currents mediating oocyte maturation events. Reproductive Biology and Endocrinology, 4(1), 1–9.

Valiente, A.G., Juanes, F. & Garcia-Vazquez, E. (2005). Reproductive strategies explain genetic diversity in Atlantic Salmon, Salmo salar. Environmental Biology of Fishes, 74(3), 323–334.

Verta, J. P., Debes, P. V., Piavchenko, N., Ruokolainen, A., Ovaskainen, O., Moustakas-Verho, J. E., Tillanen, S., Parre, N., Aykanat, T., Erkinaro, J. & Primmer, C. R. (2020) Cis-regulatory differences in isoform expression associate with life history strategy variation in Atlantic salmon. PLoS Genetics, 16(9), e1009055.

Villoutreix, R., de Carvalho, C. F., Soria-Carrasco, V., Lindtke, D., De-la-Mora, M., Muschick, M., Feder, J. L., Parchman, T. L., Gompert, Z. & Nosil, P. (2020) Large-scale mutation in the evolution of a gene complex for cryptic coloration. Science, 369(6502), 460–466.

Waters, C. D., Clemento, A., Aykanat, T., Garza, J. C., Naish, K. A., Narum, S., & Primmer, C. R. (2021) Heterogeneous genetic basis of age at maturity in salmonid fishes. Molecular Ecology. 2021 Mar;30(6):1435–56.

Weir, B.S., & Cockerham, C.C. (1984). Estimating F-statistics for the analysis of population structure. Evolution, 38, 1358–1370.

Wen, S., Ai, W., Alim, Z., Boehm, U. (2010). Embryonic gonadotropin-releasing hormone signaling is necessary for maturation of the male reproductive axis. Proceedings of the National Academy of Sciences. 107(37), 16372–16377.

Wringe, B. F., Jeffery, N. W., Stanley, R. R., Hamilton, L. C., Anderson, E. C., Fleming, I. A., Grant, C., Dempson, J. B., Veinott, G., Duffy, S. J. & Bradbury, I. R. (2018). Extensive hybridization following a large escape of domesticated Atlantic salmon in the Northwest Atlantic. Communications Biology, 1, 108,

Yeaman, S., Gerstein, A.C., Hodgins, K. A., & Whitlock, M. C. (2018). Quantifying how constraints limit the diversity of viable routes to adaptation. PLoS Genetics, 14(10), e1007717.

Yeaman S. Local adaptation by alleles of small effect. (2015). The American Naturalist, 186(S1), S74–S89.

Zheng, J., Li, Y., Abecasis, G. R., & Scheet, P. A. (2011) Comparison of approaches to account for uncertainty in analysis of imputed genotypes. Genetic Epidemiology, 35(2), 102–110.

Zueva, K. J., Lumme, J., Veselov, A. E., Primmer C. R, & Pritchard, V. L. (2021). Population genomics reveals repeated signals of adaptive divergence in the Atlantic salmon of north-eastern Europe. Journal of Evolutionary Biology, 4(6), 866–878.

