## Supplementary_Figures for "Parallel genomic basis of age at maturity across spatial scales in Atlantic Salmon"

**Supplementary Figure 1. Population structure inferred from principal component (PC) analysis in *pcadapt* of individuals genotyped on the SNP array, coloured by grilse proportion in**
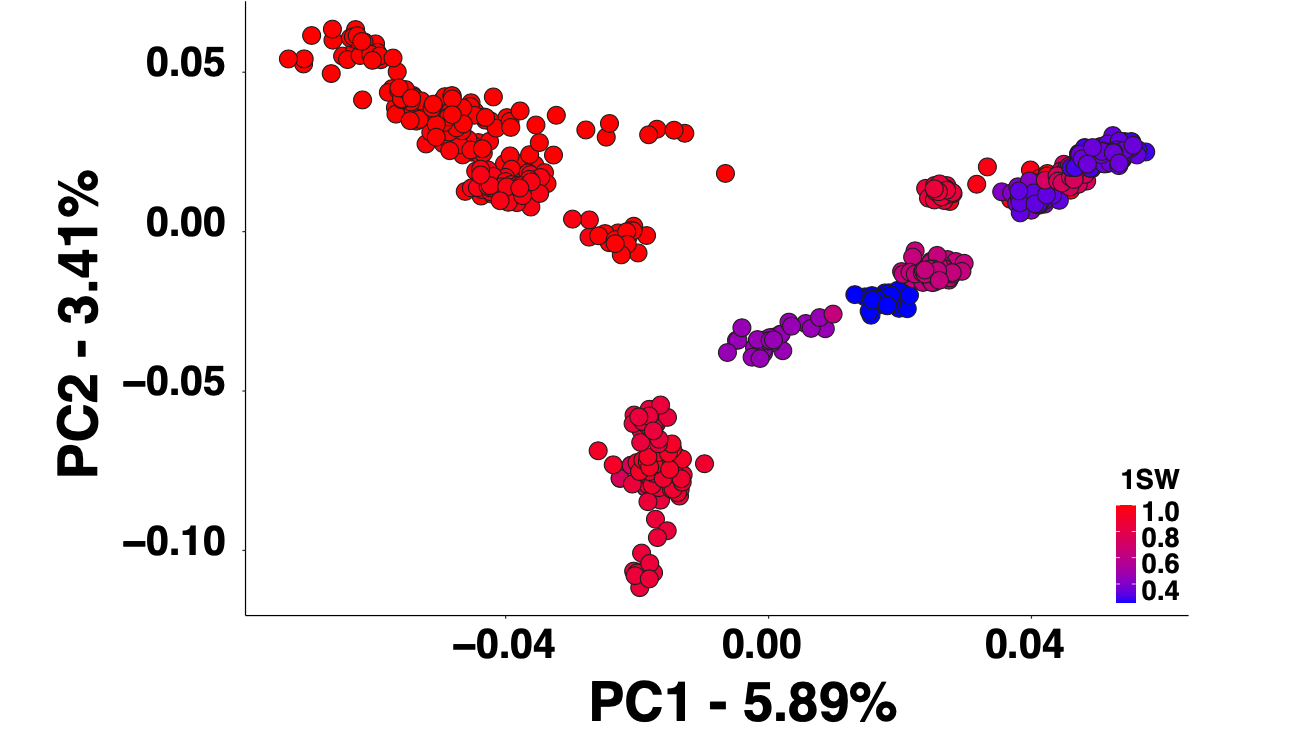
 **region of origin.**

**Supplementary Figure 2 Scree plot of variation explained by PCs at K = 10 from PCA of individuals genotyped on the SNP array using *pcadapt.***

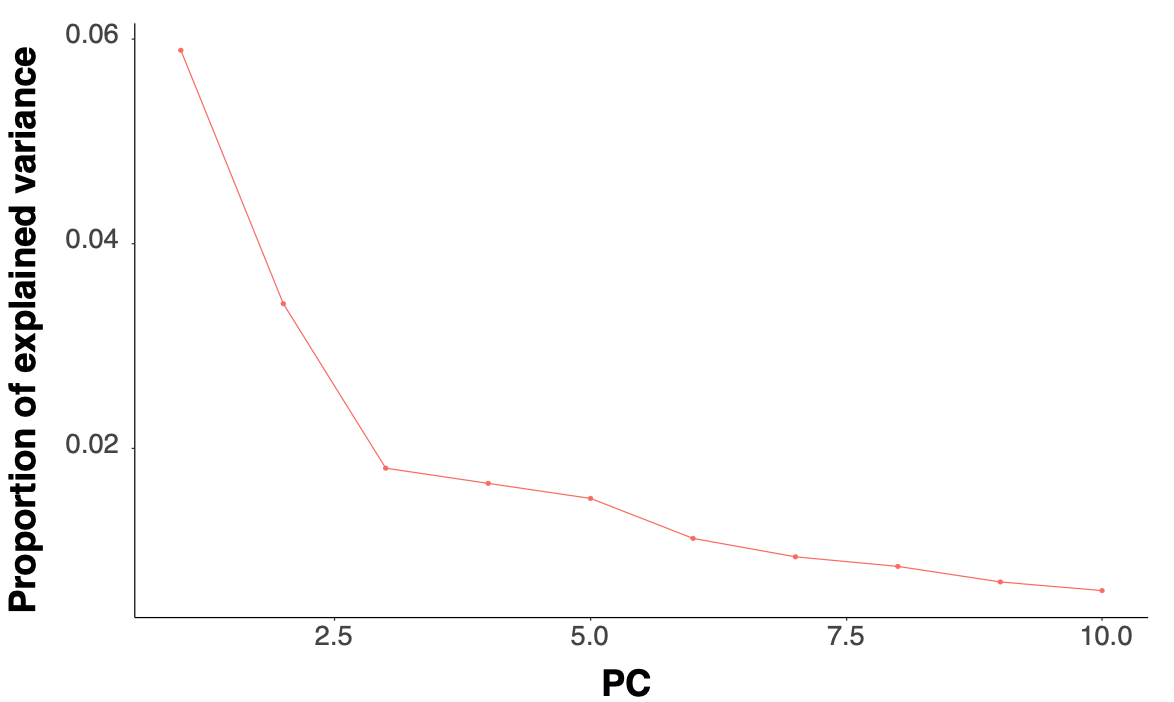

**Supplementary Figure 3. Receiver operating characteristic (ROC) curve comparing sensitivity and specificity of random forest prediction of sea age from likelihood adjusted genotype dosages at the top 500 sea age associated SNPs genotyped in 582 individuals.**

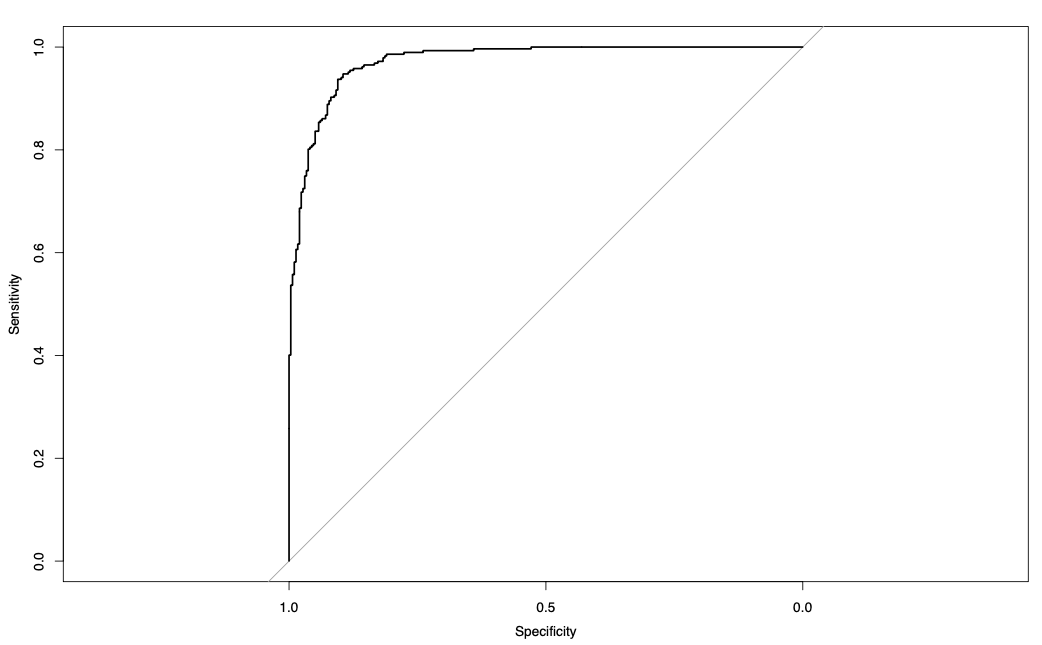
